# Crouching tiger, hidden dangers: Avian fatality rates reduced by red-blade patterning at a species-rich African wind farm

**DOI:** 10.64898/2026.05.04.722424

**Authors:** Robert E Simmons, Marlei Martins, Francisco Cervantes

## Abstract

Avian collision rates are certain to rise as renewable energy industries roll out wind and solar farms to reduce fossil fuel impacts in biologically diverse areas of the world. Technological solutions are often sought to decrease mortality rates, but for developing nations automated shut downs are expensive, and alternatives required. A promising route is to increase blade visibility to birds using high contrast colours. Despite the success of the solid black-blade experiment in Norway only one other black-blade field-study in the Netherlands has explored this possibility, with no significant results. We tested the use of colour-patterned blades at a species-rich, 37-turbine, wind facility in Hopefield, South Africa. Two broad “signal red” stripes were applied to a single blade at four high-fatality turbines, in 2023 by Umoya Energy. Avian fatality rates were compared before and after painting using the Before-After-Control-Impact (BACI) approach. Seventy-five fatalities of 23 species of raptors, passerines and wetland species over 24 months were compared for the same 20 turbines after patterning with two sets of controls: (i) their four nearest neighbours (NN) and (ii) all 16 controls (AC). Over 32 months 25 fatalities were recorded, 23 occurred at the controls and only two at the patterned turbines. Testing with Bayesian Generalized Linear Models (BGLMs) revealed a median 83% reduction in fatalities at the patterned blades for both the NN turbines (credible intervals 14% - 98%) and the AC comparisons (30% - 97%). Bayes Factors (BF) revealed strong statistical support for NN (BF = 49.9) and AC comparisons (BF = 159). There was little evidence that birds avoiding patterned turbines increased fatalities at the neighbouring turbines as there was a small median 15% increase in fatality rates when NN controls were compared with other controls, and weak statistical support (BF = 0.15). Among 14 raptor species recorded on site, 10 species have suffered fatalities. Of seven individuals killed prior to treatment at the four patterned blades, only one was killed post-treatment suggesting blade patterning is equally effective at reducing raptor fatalities. Our results show that patterned blades had a high probability (83%) of reducing fatalities with strong statistical support despite the small samples. This supports the Norway experiment in a high diversity African setting, but with red patterns not a solid black design. The strong effect of red stripes may arise from both the high contrast it provides and the possible warning effect that red may elicit. We call for additional experiments to differentiate the effect of patterns and colours for the optimal design to reduce avian-turbine collisions.

## Introduction

The use of renewable energy alternatives to reduce the world’s reliance on fossil fuels is well established in Europe and North America (Milek et al. 2022, World Resources Institute 2023). But the solution simultaneously births the green-green dilemma (Straka et al. 2020); that is, the very technology that reduces fossil fuel dependency results in a secondary green dilemma – flying vertebrates are negatively impacted (Smallwood 2013, Arnette et al. 2016, Thaxter et al. 2017, Perold et al. 2020). Fatalities arise for birds due to their apparent inability to detect moving blades and avoid them, and paradoxically this is especially so for raptorial birds despite, their legendary vision (Mitkus et al. 2018, Potier et al. 2020). Raptors not only feature prominently in fatalities around the globe (Smallwood 2003, Thaxter et al. 2017, Watson et al. 2018, Santander et al. 2026) but they are often long-lived species that are unlikely to maintain viable populations in the face of additional sources of anthropogenic mortality (Carrete et al. 2009, Kruger et al. 2015, Cervantes et al. 2022).

This increased pressure on raptors is especially pronounced in Africa, a continent rich in 108 species of raptor (Clarke and Davies 2018), a taxon already under severe threat from anthropogenic impacts (Simmons et al. 2004, Ogada et al. 2015, Cervantes et al. 2022, Shaw et al. 2023). As a developing region with endemic poverty levels but abundant resources, Africa needs to develop renewables to power its poorer nations. However, as more areas of low environmental risk are taken for wind farming, developers will move to more risky areas (Santangeli et al. 2016), triggering the need for additional or more effective forms of mitigation (Marques et al 2014, Bennun et al. 2021, Katzner et al. 2025). This comes at a high cost for automated shut down on demand technologies in biologically diverse nations where hundreds of species are at risk of collision (Perold et al. 2020, Ralston-Paton 2025, Vargas-Jimenez and Rico 2025, Santander et al. 2026, Roy et al. 2026). This necessitates alternative solutions to reducing collision impacts, of which the most innovative is to mark (pattern) the blades with high contrast colours to make them more visible to at-risk birds (McIsaac 2001, Hodos 2003, Martin and Banks 2023, Blary et al. 2024).

Promising first field test results emerged from Norway where White-tailed Eagles *Haliaeetus albicilla* were regularly killed at the Smøla Wind farm’s white-painted turbines prior to the addition of a solid black design to one blade of four turbines (May et al. 2020). This was deemed the best design to break up “motion smear” - the inability of the retina to detect movement for fast spinning objects – in lab experiments (Hodos 2003). Fatality rates dropped by 72% for all birds (May et al. 2020) at the painted turbines. The efficacy of the black-blade approach in other environments has recently been thrown into doubt by a replica field experiment on the Dutch coast where seven experimental turbines showed no significant reduction in avian fatalities relative to un-patterned controls (Klop et al. 2025). An alternative design suggested by McIsaac’s lab experiments (2001) was to pattern blades with two broad transverse black stripes, and this is the basis of our approach.

The aim of this paper is to describe the first trial of colour-patterned blades as a mitigation. Using a Before-After-Control-Impact (BACI) design across 32 months of post-treatment monitoring, we tested the hypothesis that two broad Signal Red stripes on a single blade would reduce avian fatalities at treated turbines relative to un-patterned controls. Red was used in preference to black due to constraints imposed by South African civil aviation authorities on the use of black on tall structures. Because birds avoiding patterned turbines might veer towards neighbouring control turbines, we simultaneously tested for increased fatalities at nearest neighbours. Given the prominence of raptors in wind-farm fatalities globally, and the presence of 14 raptor species at Hopefield, we simultaneously assessed whether raptors reflected any treatment effect. This is particularly important for one threatened species the Black Harriers *Circus maurus* whose population viability is highly sensitive to additional anthropogenic impacts.

### Study Area

The Umoya Wind farm near Hopefield (S33° 05’ 43” E18° 23’ 32”) in the Western Cape of South Africa (Figure 1) is situated in a mix of farmland and native *Endangered* Hopefield Sand Fynbos (Mucina and Rutherford 2011) in the mega-diverse Cape Floral Kingdom (Cowling and Richardson 1995, Curtis-Scott et al. 2020), about 2 hours north of Cape Town. Most precipitation falls in winter with a mean annual precipitation of 326 mm and mean maximum and minimum temperatures of 28.3° C and 7.1° C respectively. Foggy days occur often enough (110 days per year) for the harvesting of fog to be researched locally (Olivier 2004). Daylight hours for Cape Town range from 14.4 h at the height of summer to 9.54 hours in winter (Sunrise and sunset in Cape Town (timeanddate.no).

**Figure 1:**
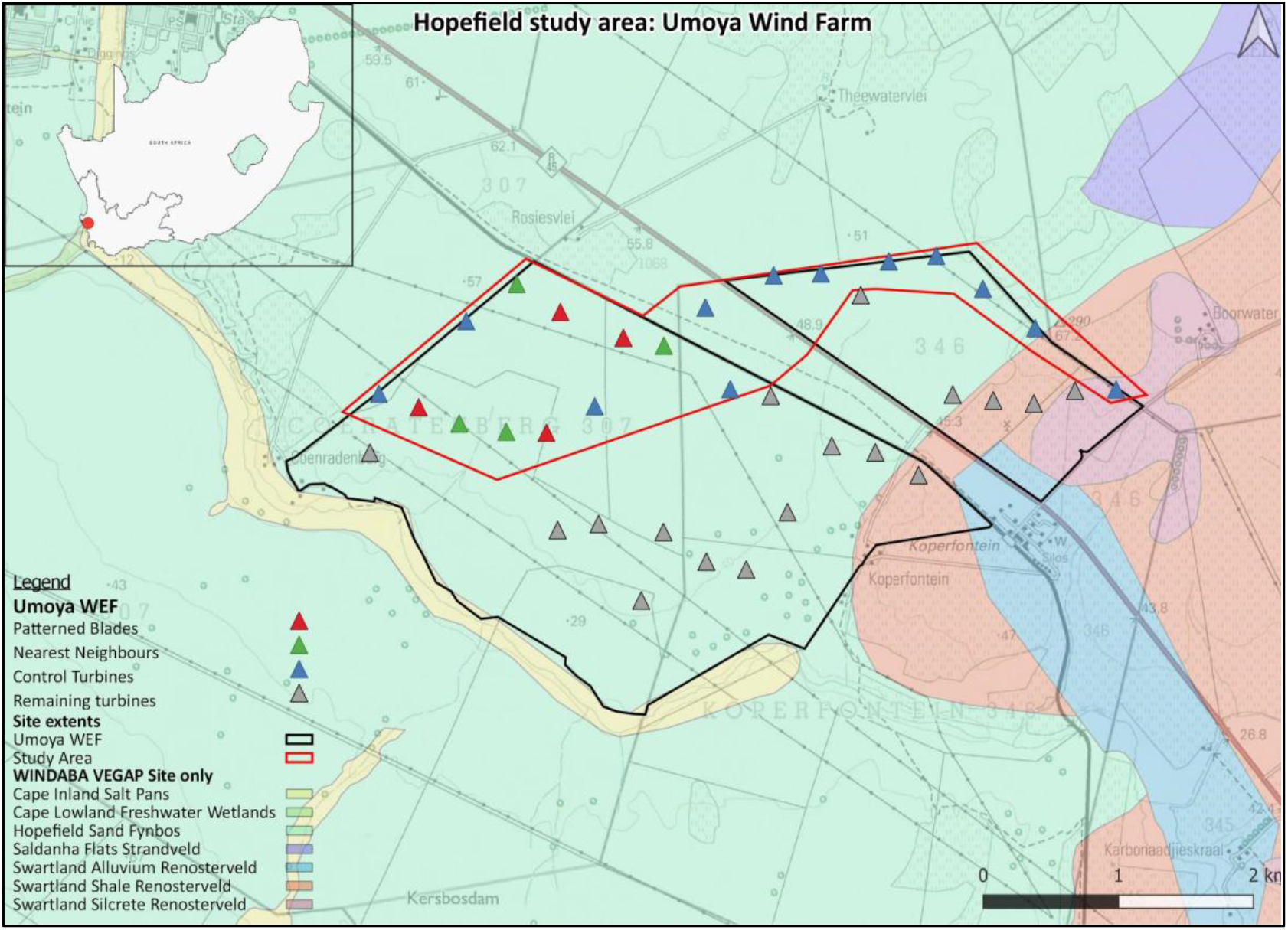
Umoya wind farm indicating locations of all turbines (triangles) and the four experimental (red triangles) and control turbines (green: nearest neighbour, and blue: all other controls), near Hopefield, South Africa. Turbines not used in the BACI experiment are shown as grey triangles. Inset: red dot shows location of Hopefield in South Africa.

**Figure 2:**
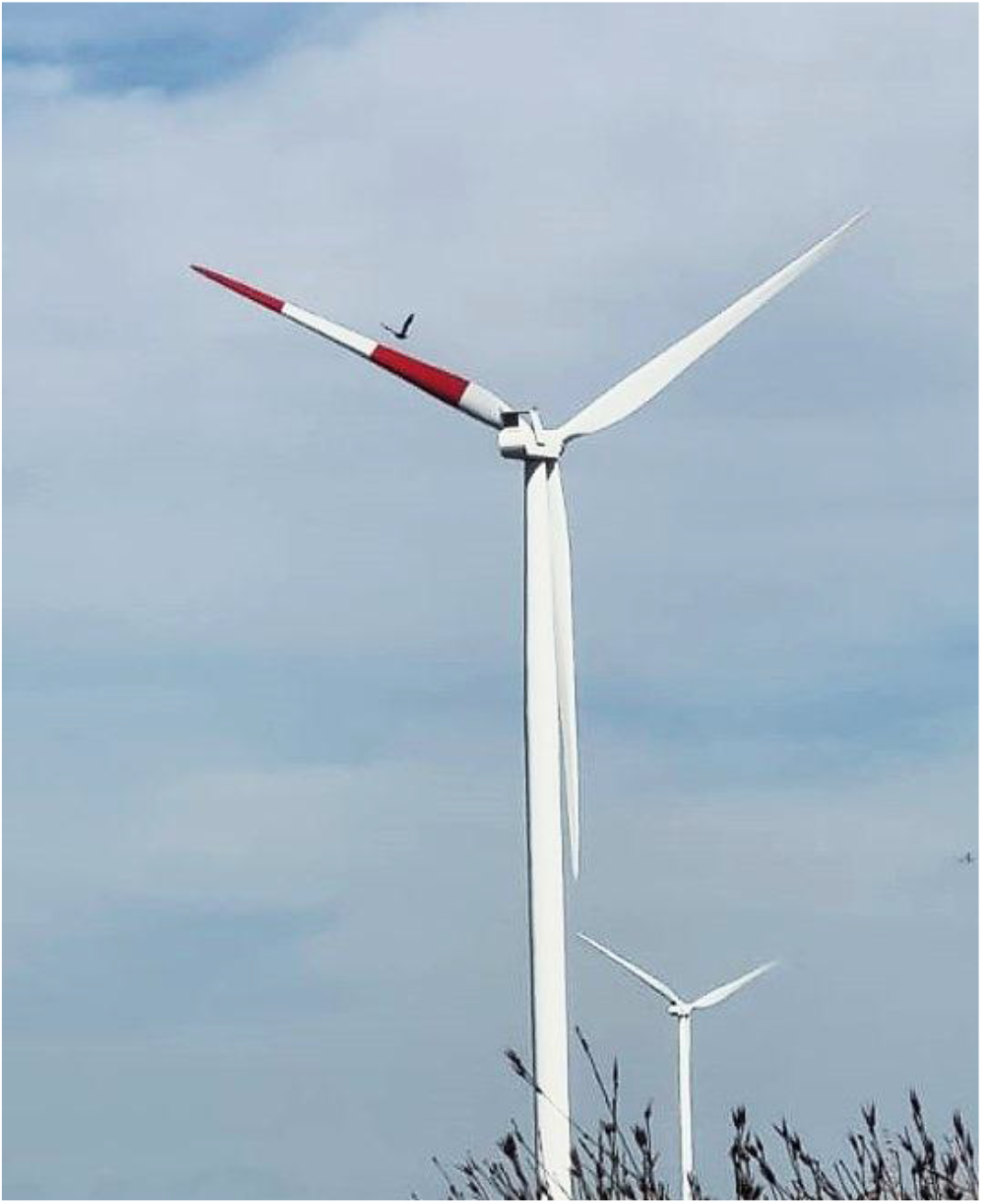
An adult Jackal Buzzard veers away from an Umoya Wind farm turbine patterned with two broad Signal Red stripes in October 2023.

The wind farm comprises 37 Vestas wind turbine generators of 1.8 MW each and an installed capacity of 66.6 MW. The hub height (ground to nacelle) is 95m with 50-m, Pure-white (RAL 9010) blades. Umoya Energy (Pty) Ltd was the first commercial farm in South Africa to receive Environmental Authorisation (September 2009) and has been wind farming since 2014.

The avian community recorded at the Hopefield wind farm comprises at least 97 species of which 15 are collision-prone (Arcus 2016), that is, occurring within the top 100 priority species susceptible to wind farm collision in South Africa (Ralston et al. 2017). Of these, 14 raptorial (Accipitridae) species have been recorded at Hopefield, including globally or regionally *Endangered* Black Harriers (breeding), and Martial Eagles *Polemaetus bellicosus* (breeding) and *Vulnerable* African Marsh Harriers *C. ranivorus* (Lee et al. 2025). South Africa’s most collision-prone species, the Jackal Buzzard *Buteo rufofuscus* (Ralston-Paton 2025), also occurs and breeds at the site. Because raptors formed a high proportion of the fatalities in early monitoring, blade patterning was suggested as a cost-effective mitigation to reduce that.

## Methods

Lengthy discussions with the South African Civil Aviation Authority (SACAA) lead to an approval in August 2022 for Umoya Energy Wind farm to pattern four blades. This was necessary to change the blade colour from Pure White (RAL 9010, reflectance 85%) to one with two broad red stripes at four operational turbines. Signal Red (RAL 3001, reflectance of 8%) is permitted on tall structures under SACAA regulations but black is not, hence the evolution of red stripes. Blade painting was undertaken over 3 months (January to March 2023), *in situ* from a raised platform, and two coats applied. All four patterned turbines were in operation by April 2023 and that coincided with the start of the 32 months’ of weekly carcass searching by the Umoya team reported here.

The study was undertaken in three phases: (i) an initial 12-month carcass survey of all 37 un-patterned turbines by Arcus Environmental in 2015-2016 (Arcus 2016), followed by (ii) a 12-month carcass survey in 2020-2021 overseen by Birds & Bats Unlimited (BBU 2021) and then (iii) the experimental phase beginning in April 2023 (after blade patterning) for 32 months. Throughout, we followed the same methods of carcass searching, within an 80m radius of the tower, walked by two observers 6 m apart. This was undertaken weekly to cover all 37 turbines in the first two before-treatment surveys (by two pairs of observers). In 2023, once the four turbines were patterned (January-March 2023), carcass searching was resumed at 20 of the 37 turbines, employing the same methods, with one team of two people. This allowed teams to focus on a smaller study site and make up for any days lost to inclement weather when not allowed on the wind farm. This included the four patterned turbines and 16 control turbines, 54% of the wind farm. All turbines were surveyed equally unless they went offline; on one occasions carcasses at the nearest operational turbine were counted until the target turbine came online again.

For conservation purposes, we chose four of the nine turbines that had previously killed the most birds or red data species, for treatment. We assigned the four nearest neighbours as controls regardless of the number of fatalities they had been responsible for. We recommended two broad Signal Red stripes, each 25% the length of the blade (i.e., 21.25m) starting at the blade tip and separated by an equal space of 21.25 m of Pure White paint (Figure 1). This pattern was based on McIsaac (2001) whose laboratory research showed that American Kestrels *Falco sparverius* (and student volunteers) found this pattern on moving turbine blades more conspicuous than other designs, including plain white, plain grey or blades with multiple cross-sectional (zebra) stripes or two longitudinal stripes.

### Statistical analysis

We employed the powerful BACI (Before-After-Control-Impact) analysis (Smith 2002) because carcass searching had been undertaken at all 37 turbines for 2 years before patterning occurred. That is, the (four) turbines to be patterned were sampled for carcasses both before and after painting, as were other (un-patterned) turbines. Two sets of un-patterned turbines were chosen as controls: (i) the four nearest neighbours (NN) immediately adjacent to the patterned turbines and (ii) the 12 Other Control (OC) turbines within the same region. We chose the NN because they were more likely to be matched for habitat, wind resource and aerial traffic as the patterned blades and thus made ideal controls. We also compared fatalities at the 4 NN turbines with the 12 OC turbines to determine if birds avoiding the patterned blades were diverted into the NN turbines (May et al. 2020). No other mitigations such as observer-led, automated shut downs or curtailments (Smallie et al 2025) were in operation during these experiments, simplifying the BACI comparisons. A total of 20 turbines were surveyed in each site visit (4 patterned, 4 NN, 12 OC).

We used a Bayesian Generalised Linear Model (BGLM) to determine if, and by how much, patterned blades reduced fatalities. We employed a Poisson generalized linear model, using number of fatalities per turbine and year as the response variable. As explanatory variables we used indicator variables for the periods Before-After treatment, to differentiate turbines into Treatment-Control, and most importantly for the BACI analysis, the interaction between the two. The latter tells us about the differential effect of the Before-After periods in relation to the Treatment-Control turbines. We tested for an effect of year and a random intercept for each turbine. Thus, we fitted three different models: 1) a simple BACI with no other covariates, 2) a BACI with an effect of year, and 3) a BACI with an effect of year and a random intercept for individual turbines. We compared these models using Pareto Smoothed Importance Sampling leave-one-out cross validation (PSIS-LOO, Vehtari et al, 2017). With the most parsimonious model, we analysed the interaction term as the reduction factor in fatalities at the patterned turbines in relation to the controls, after treatment. We also performed a BACI contrast whereby we computed *Δ* = (*AI* − *BI*) − (*AC* − *BC*), which tell us the difference in fatalities reduction between patterned turbines and controls, after treatment. Finally, to determine the statistical support for the hypothesis of a reduction in fatalities at the patterned turbines we computed the Bayes Factor (BF) of *Δ* < 0, *BF*=*Pr* (*Δ* < 0)/*Pr*(*Δ* ≥ 0). We considered BF between 3 and 10 to provide moderate statistical support and a BF > 10 to provide strong statistical support (Kass and Raftery 1995).

We conducted our analyses in R using the package cmdstanr to use the Bayesian software Stan. We ran four chains with 4000 samples each using the No-U-turn Hamiltonian Monte Carlo algorithm and discarded the first 2000 as adaptation. We assessed model convergence with the Gelman-Rubin statistic, which compares the intra-chain variation with the inter-chain variation, with values between 0.9 and 1.1 indicating convergence (Gelman and Rubin, 1992). PSIS-LOO model comparison was implemented using the loo package (Vehtari et al., 2024).

## Results

### Pre-patterning fatalities

A total of 75 bird carcass of 23 species were recorded under all 37 turbines over 24 months before patterning: a low fatality rate (unadjusted) of 1.01 birds/turbine/yr. In the first year, 40 avian fatalities (of 12 species) were recorded at the 37 turbines (2014-2015) and 35 birds (of 15 species) in the second year of 2020-2021 (Table 1). Of these 33% were raptorial species (Table 1), with the remainder comprising passerines (22%), gamebirds (20%), doves (12%) and wetland birds (1%).

**Table 1.** Avian fatalities in two years’ pre-treatment monitoring at all 37 turbines at the Umoya, Hopefield wind farm by Arcus Consulting (2014-2015) and Birds & Bats Unlimited (2020-2021). Highly collision-prone species are shown in bold. Cape Spurfowl were omitted from further analyses as tower impacts. Full species totals given in Suppl Table 1.

| Species | Collision fatalities |  | Turbine identity |  |
| --- | --- | --- | --- | --- |
|  | 2014-15 | 2020-21 | 2014-15 | 2020-21 |
| African Spoonbill |  | 1 |  | T22 |
| Ant-eating Chat |  | 1 |  | T13 |
| Barn Owl |  | 3 |  | T6, T31, T21 |
| **Black Harrier** | **1** | - | **T31** |  |
| **Black-shouldered Kite** | **5** | - | **T17,T17,T23,T32,T37** |  |
| Bokmakierie | 7 | 2 | T3,T3,T7,T11,T19,T20,T28 | T22 T37 |
| **Booted Eagle** | **3** |  | **T28,T30,T36** |  |
| Buzzard species | 2 |  | T18, **T28** |  |
| Cape Bulbul | 1 |  | T14 |  |
| Cape Sparrow | 1 |  | T1 |  |
| Cape Spurfowl | 7 | 7 | T25, T28, T28, T28, T28, T34, T34 | T21,T28,T30,T30,T32, T13, T22 |
| Cape Turtle Dove | 1 |  | T28 |  |
| **Jackal Buzzard** | **4** |  | **T1, T1, T35, T36** |  |
| Helmeted Guineafowl |  | 1 |  | T27 |
| Large-billed Lark |  | 1 |  | T5 |
| Lark species |  | 1 |  | T2 |
| Little Swift | 2 |  | T7,T35 |  |
| Racing Pigeon |  | 1 |  | T31 |
| Red-eyed Dove |  | 1 |  | T15 |
| Swift |  | 1 |  | T14 |
| Rock Kestrel | 4 | 2 | T25,T29,T34, T35 | T27, T5 |
| Speckled Pigeon | 4 | 2 | T25, T29, T34, T35 | T12, T23 |
| White-backed Mousebird |  | 1 |  | T25 |
| **Yellow-billed Kite** |  | **1** |  | T7 |
| Unknown passerine |  | 3 |  | T20 T13, T32 |
| Unknown bird |  | 6 |  | T30, T34, T6, T29, T23, T2 |
| **SUMMARY: 23 species** | **40 (13)** | **35 (1)** | **Four turbines killed 3 or more birds (T28, T31, T34, T35); these were chosen for patterning** |  |

Fatalities recorded at the subset of 20 turbines totalled 51 birds of which 18 (35%) were raptors before patterning (STable1).

Evidence from all years suggested that Cape Spurfowl *Pternistis capensis* were tower-collisions, not blade strikes, because they were found within a few metres of the towers and showed no traumatic injuries. From the 20 turbines and the total 51 fatalities recorded before treatment, 12 were tower-collisions and removed from further analyses.

Nine of the 37 turbines accounted for 50% of the recorded fatalities (37 of 75) and four of the nine turbines (T28, T31, T34 and T35) were responsible for 25% of all 75 fatalities: these were chosen for patterning. With tower-collisions removed the four turbines selected for treatment were responsible for 24% of the 55 fatalities on site.

### Number and species composition of fatalities before and after patterning

Prior to patterning 75 individuals of 22 species of bird were killed by the 37 unmarked turbines over a 24-month period. These comprised 25 individual raptors (eight species), 17 individual passerines (six species), nine individual doves (four species), 15 individual gamebirds (two species), three individual hirundines (one species) and one wetland bird (1 species). Six unidentified birds were also recovered beneath the turbines (Table 1).

After patterning, the 20 turbines killed 42 individuals of 15 species over a 32-month period. These comprised nine individual raptors (five species), 11 individual passerines (four species), 21 individual game birds (four species), and two wetland birds (two species).

At the four experimental turbines, 20 individuals were killed prior to patterning comprising seven individuals raptors (six species), seven individual gamebirds (one species), one individual passerine (one species), three individual doves (three species), one hirundine (one species), one individual wetland bird (one species) and one unidentified individual (Suppl Table 1). The tower impacts were removed from further analyses. After patterning only two individual birds of two species were killed in 32 months: one *Milvus* kite and one shrike.

### Post-patterning fatalities: treatment vs nearest neighbours (NN)

Here we compare the reduced fatalities at the experimental turbines (from 13 individuals before to 2 individuals after treatment) with fatalities recorded simultaneously at the NN turbines. In 24 months before treatment seven fatalities were recorded, identical to those recorded in the 32 months after-treatment at the nearest neighbour controls (Figure 3).

**Figure 3:**
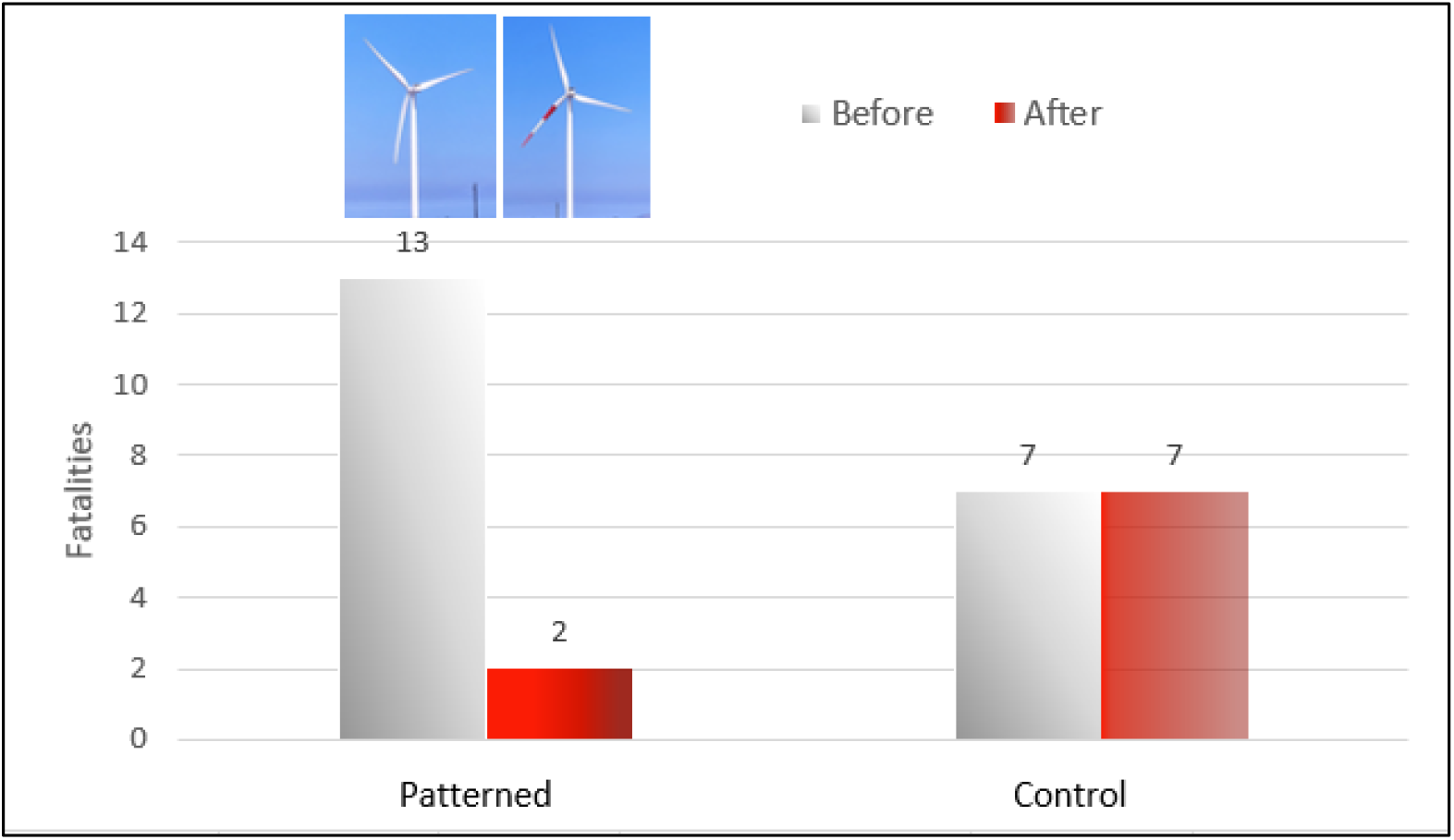
Influence of blade patterning on avian fatalities for the 4 patterned blades vs the 4 nearest neighbours (Control) after 32 months. The left-hand pair of bars indicate fatalities for the experimental turbines before and after treatment. The right-hand bars show the fatalities for the control turbines before and after treatment.

Model comparison with PSIS-LOO selected the simple BACI setup with no covariates and no random effects as the most parsimonious model (Table 2). This model (Table 3) indicates a median reduction in fatalities at the patterned turbines 83% greater than that at the NN controls (95% credible interval: 14% - 98% greater reduction). The BACI contrast estimates a median reduction of 1.2 more fatalities per turbine at the patterned turbines than at the NN controls (95% credible interval: 0.06 - 2.6 greater reduction) with strong statistical support for a greater reduction at patterned turbines (Bayes Factor = 49.9).

**Table 2.** Model comparisons (using PSIS-LOO test) between three model types: mod1) a simple BACI with no other covariates; mod2) a BACI with an effect of year; and mod3) a BACI with an effect of year and a random intercept for individual turbines. For each model we show comparisons of: 1) treatment vs nearest neighbours (Treat-NN), 2) treatment vs all controls (Treat-ALL) and 3) nearest neighbours vs all controls (NN-ALL). We provide the estimated log predictive density (elpd), together with its standard error (SE).

| comparison | Model | elpd | SE |
| --- | --- | --- | --- |
| Treat-NN | mod1 | -94.62 | 6.30 |
| Treat-NN | mod2 | -95.32 | 6.50 |
| Treat-NN | mod3 | -94.84 | 5.96 |
| Treat-ALL | mod1 | -40.47 | 4.39 |
| Treat-ALL | mod2 | -41.50 | 4.64 |
| Treat-ALL | mod3 | -40.62 | 4.14 |
| NN-ALL | mod1 | -40.47 | 4.39 |
| NN-ALL | mod2 | -41.50 | 4.64 |
| NN-ALL | mod3 | -40.62 | 4.14 |

**Table 3.** Generalized linear mixed model summary for BACI comparison of patterned blades with nearest neighbour turbines. We show for each factor in the model: the mean, standard deviation (sd), the 5% and 95% percentiles (q5, q95), the Gelman-Rubin statistic (rhat) and the effective sample size for the posterior distribution (ESS).

| variable | mean | Sd | q5 | q95 | rhat | ESS |
| --- | --- | --- | --- | --- | --- | --- |
| Intercept | -0.160 | 0.367 | -0.803 | 0.402 | 1.001 | 2559.422 |
| After | 0.587 | 0.458 | -0.141 | 1.373 | 1.001 | 2692.490 |
| Impact | -0.084 | 0.528 | -0.953 | 0.768 | 1.000 | 2585.703 |
| After*Impact | -1.840 | 0.905 | -3.375 | -0.427 | 1.001 | 3228.917 |

### Post-patterning fatalities: treatment vs all controls (AC)

Here we compare the reduced fatalities at the patterned blades (13 fatalities to 2) with all controls. In 24 months before treatment 26 fatalities were recorded, and in the 32 months after treatment 23 fatalities were simultaneously recorded at all controls (Figure 4).

**Figure 4:**
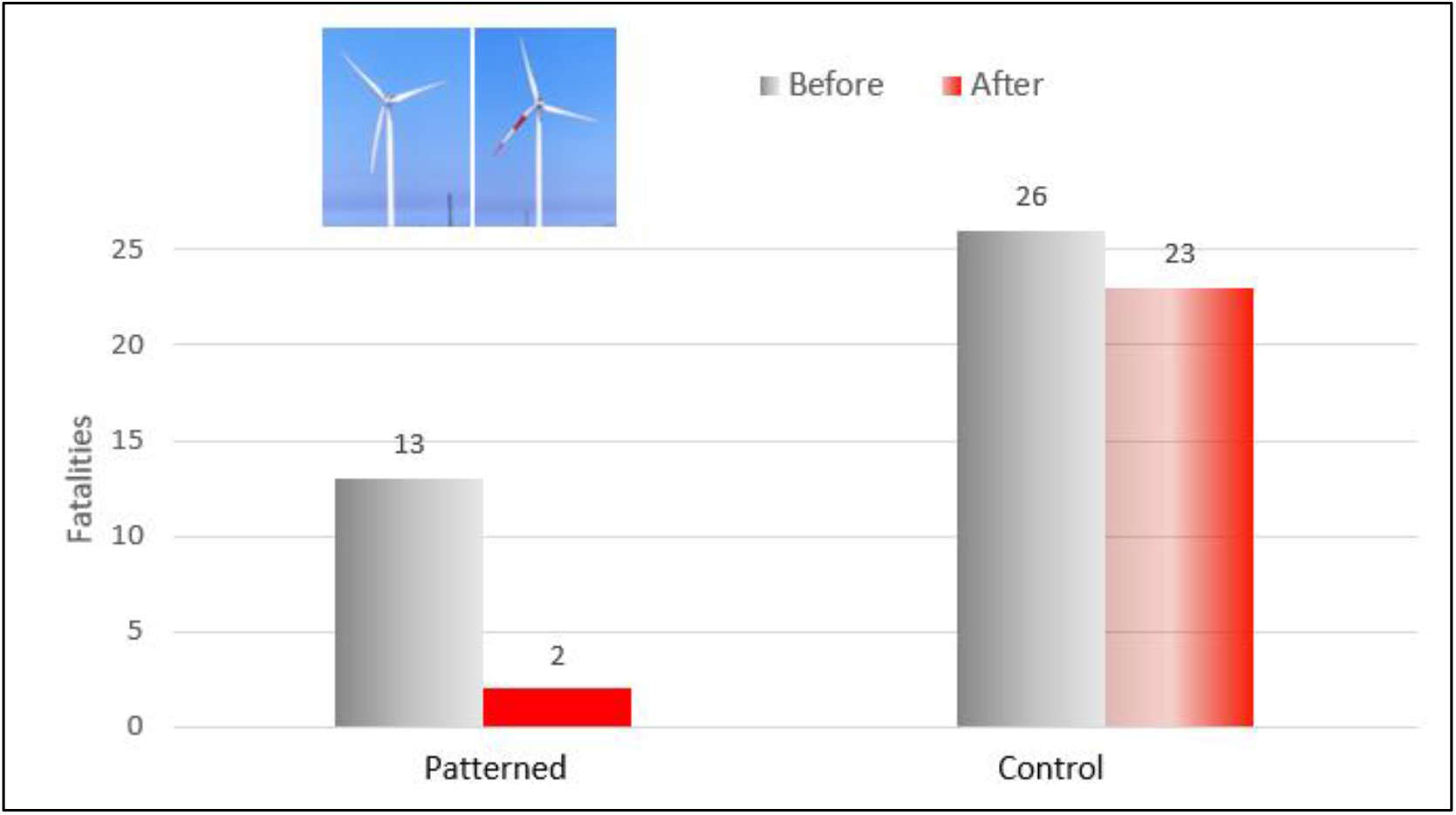
Influence of blade patterning on avian fatalities for the 4 patterned turbines vs all 16 Controls after 32 months. The left-hand pair of bars indicate fatalities for the experimental turbines before and after treatment. The right-hand bars show the fatalities for the control turbines before and after blade patterning.

The statistical analysis showed very similar results to those obtained with the NN controls. Model comparison with PSIS-LOO (Table 2) again selected the simple BACI setup with no covariates and no random effects as the most parsimonious model (Table 4). This model indicates a median reduction in fatalities at the patterned turbines 83% greater than that at the NN controls (95% credible interval: 30% - 97% greater reduction). The BACI contrast estimates a median reduction of 1.2 more fatalities per turbine at the patterned turbines than at the NN controls (95% credible interval: 0.26 - 2.3 greater reduction) with a very strong statistical support for a greater reduction at patterned turbines (Bayes Factor = 159).

**Table 4.** Generalized linear mixed model summary for BACI comparison of patterned blades with all 16 turbines as controls. We show for each factor in the model: the mean, standard deviation (sd), the 5% and 95% percentiles (q5, q95), the Gelman-Rubin statistic (rhat) and the effective sample size for the posterior distribution (ESS).

| comparison | mean | sd | q5 | q95 | rhat | ESS |
| --- | --- | --- | --- | --- | --- | --- |
| Intercept | -0.210 | 0.191 | -0.535 | 0.104 | 1.001 | 3638.870 |
| After | 0.634 | 0.347 | 0.054 | 1.194 | 1.001 | 4145.104 |
| Impact | -0.148 | 0.274 | -0.604 | 0.300 | 1.001 | 4056.862 |
| After*Impact | -1.806 | 0.819 | -3.246 | -0.554 | 1.001 | 5121.080 |

**Table 5.** Generalized linear mixed model summary for BACI comparison of NN turbines with the remaining 12 control turbines. We show for each factor in the model: the mean, standard deviation (sd), the 5% and 95% percentiles (q5, q95), the Gelman-Rubin statistic (rhat) and the effective sample size for the posterior distribution (ESS).

| Comparison | Mean | sd | q5 | q95 | rhat | ESS |
| --- | --- | --- | --- | --- | --- | --- |
| Intercept | -0.263 | 0.229 | -0.649 | 0.106 | 1.001 | 3350.151 |
| After | 0.075 | 0.446 | -0.696 | 0.775 | 1.001 | 3430.039 |
| Impact | -0.169 | 0.339 | -0.729 | 0.382 | 1.001 | 3236.733 |
| After*Impact | 0.146 | 0.635 | -0.91 | 1.197 | 1 | 3320.823 |

### Do the patterned blades increase fatalities at their nearest neighbours?

The decrease in fatalities at the patterned turbines may arise because approaching birds veer away from the patterned turbines and into the un-marked nearest neighbours (NN). We tested this possibility by comparing fatalities before and after patterning for the four NN turbines relative to the remaining 12 OC turbines. We would expect increased fatalities at the nearest neighbour controls after treatment, but no such increase was apparent (Fig. 5).

**Figure 5:**
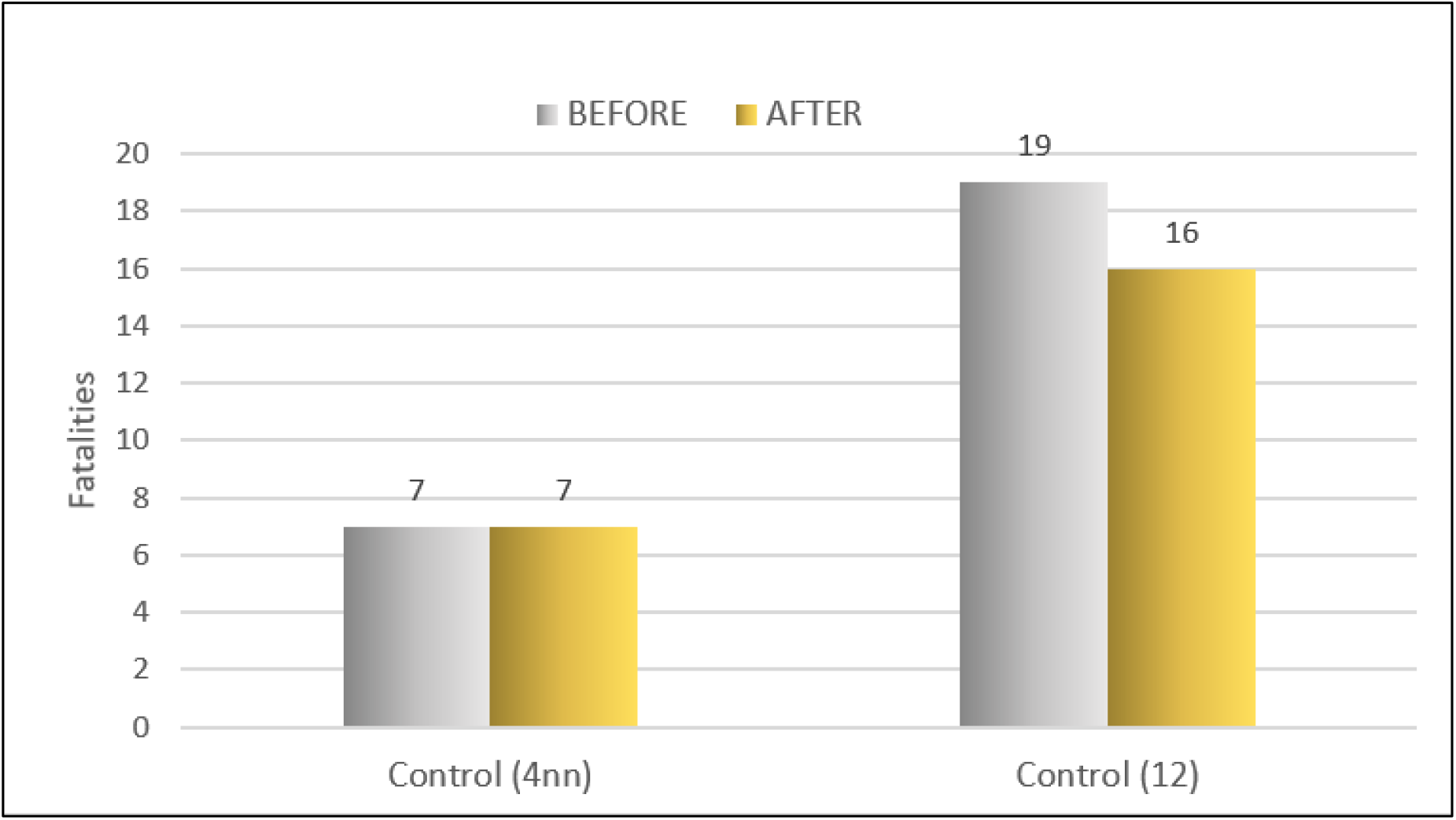
Avian fatalities at nearest neighbours (nn) before and after painting (left hand bars) relative to the other (12) controls (right hand bars) before and after painting of the four experimental turbines (not shown).

Model comparison with PSIS-LOO again selected the simple BACI setup with no covariates and no random effects as the most parsimonious model (Table 3). This model indicates a median increase in fatalities at the NN control turbines 15% greater than that at all controls (95% credible interval: 315% greater increase - 67% greater reduction). The BACI contrast estimates a median increase of 0.1 more fatalities per turbine at the NN control turbines than at all controls (95% credible interval: 1.13 greater increase - 0.9 greater reduction).

There was weak statistical support for a greater increase at NN controls turbines in relation to all controls (Bayes Factor = 0.73).

### Reduction in fatalities at patterned turbines

An alternative method of determining the magnitude of the reduction in fatalities is to calculate the fatality rates per turbine per month for all patterned and all control turbines prior to treatment and use these to calculate expected rates after treatment. At the rates found prior to patterning (13 fatalities in 24 months), we expected 14.1 fatalities at the four patterned blades in 32 months after treatment. However, only two were recorded in the same period. For the 16 control turbines, however, at the fatality rate calculated (24 fatalities in 24 months) the expected fatalities (32.3) were similar to that observed (23) after 32 months (Table 6). At these rates the patterning reduced the fatalities 88.4% for all birds, over pre-patterning rates in the 32 months of post-treatment monitoring.

**Table 6.** Avian fatalities predicted (from the rate of fatality in 24 months’ prior to the patterning of blades turbines) in relation to that observed for patterned and 16 Control turbines in 32 months post-patterning, at the Umoya wind farm at Hopefield.

|  | Fatality rate before patterning | Fatalities **expected** in 32 months | Fatalities **observed** in 32 months | Reduction in fatalities based on previous rates |
| --- | --- | --- | --- | --- |
| Patterned Turbines (n = 4) | 0.135 fatalities/T/month | 17.3 | 2 | 88.4% |
| All Control Turbines (n = 16) | 0.063 fatalities/T/month | 32.3 | 23 | 28.8% |

Among the species of birds impacted by the turbines before treatment, most were raptors (33% of 75), passerines (27%) and gamebirds (20%). At the four experimental turbines seven individual raptors of six species were killed before patterning, including an *Endangered* Black Harrier (Table 1). However, only one raptor (a young, Yellow-billed Kite *Milvus parasitus*) was killed in the 32 months following the treatment at the patterned turbines (Suppl Table 1). This was despite (i) the presence of six raptor species regularly in the farm, (ii) three pairs of collision-prone Black Harriers breeding in the facility in 2024 /2025 and (iii) eight raptor fatalities recorded at the control turbines, including a non-breeding Black Harrier. The reduction in raptor fatalities based on the rate of fatalities before treatment (0.7 raptors per turbine per month) was similar to that for all birds at 86.8%.

## Discussion

We have shown here, in the first global field trial of red-patterned turbine blades at four operational turbines in South Africa, that expected avian fatality rates decreased by 83% over 32 months. The Bayesian assessment of the BACI experiment revealed that neither year nor turbine identity (included as random effects) played a role in the reduced fatalities at these previously high fatality-rate turbines following treatment. This occurred whether the comparison was with the NN turbines or for the larger set of 16 AC turbines relative to the experimental turbines.

Given that we had only four experimental turbines, due to financial constraints on painting turbines *in situ*, it is not surprising that the credible intervals were wide for the NN comparison (14% to 98%) and slightly less so for all controls (30% to 97%). This means that there is a high degree of uncertainty associated with these median values. Nevertheless, we found strong statistical evidence for an effect, as indicated by the large Bayes Factors obtained when comparing the hypothesis of a larger reduction in fatalities at the patterned turbines against the control turbines.

These results are based on un-corrected carcass searches. Correction factors for scavenger removal and non-detection by observers in the first surveys (following the Korner-Nievergelt et al. 2015 methods) indicates that only about half (52.6%) of all bird carcasses were located (Arcus 2016). Given that this bias should influence carcass detection at both experimental and control turbines in the same way, it should not affect the overall results. However, it approximately doubles (1.9x) the number of fatalities recorded at the Umoya wind farm. Adjusted fatality rates thus rise to ~1.93 birds/turbine/year, still less than half the average for South African wind farms (4.25 birds/turbine/year: Ralston-Paton 2025).

The carcass searchers reported numerous Cape Spurfowl fatalities next to the towers throughout all monitoring periods. Inspection of the carcasses revealed no traumatic injuries (i.e. missing limbs or heads). Tower impacts are reported for game birds in Norway (Stokke et al. 2020) and Spain (FC pers obs) and for multiple species in the USA adjacent to the towers (Choi et al. 2020). We removed all individuals (30 0f 94 carcasses) from the analysis because patterning the turbine blades was not expected to alter the behaviour of birds colliding with the towers. Indeed, identical numbers of tower strikes were found before (7) and after (7) treatment at the experimental turbines, supporting the idea that blade patterning makes no difference to tower strikes. However, painting tower bases black, independently of blade painting, did reduce tower fatalities in Norway (Stokke et al. 2020).

Our results support the pioneering field test in Norway (May et al. 2020) that first tested the idea of Hodos (2003) that breaking up the “motion smear” with the addition of a single black blade increases their visibility and reduces avian impacts. We tested a slight modification of this idea in that we painted one blade but used the two broad stripes (not a single continuous design) as it was the most conspicuous pattern to kestrels in the lab (McIsaac 2001). Our experiment cannot distinguish between motion smear (a single blade marked) and increasing contrast (blades with high contrast stripes) as the reason for significantly reduced impacts. However, it is more likely that the striped patterns used at Hopefield elicited the avoidance behaviour given that two lab experiments with similar transverse stripes invoked the strongest behavioural responses (McIsaac 2001, Blary et al 2026) but patterned all three blades. Patterning all three blades argues against motion smear as an explanation for their results.

### Patterns vs solid blade painting designs and colour

At Eemshaven, on the Dutch coast, Klop et al. (2025), found no significant reduction in fatalities at seven black-blade turbines with appropriate controls after 24 months of post-treatment surveys. That “Diurnal species do show a reduction in collision rates after painting the blades, but the reduction is fairly weak” (Klop et al. 2025) suggests that nocturnal migrants were not detecting the dark blades, or that the nearby port of Eemshaven provided a visually confusing backdrop for the birds. In the Hopefield experiment, testing a patterned blade design (Figure 2), rather than a solid (black) blade has provided significant results in 32 months’ of monitoring, despite a low rate of bird strikes as observed in the Smøla experiment (1.0 vs 1.1. birds/turbine/year).

This unusual Eemshaven result begs the question, what are the cues allowing some birds to detect moving or stationary blades while others, under different conditions, do not. One hypothesis is that certain species of *Gyps* vultures, and bustards (*Otis* species) may be susceptible to collisions mainly because of limits imposed by their skull morphology, limiting vision directly ahead (Martin and Shaw 2010, Martin et al. 2012). However, if this were the main reason for blade strikes of numerous raptors, the addition of high chromatic contrast colours to the blades would be expected to have little effect if the birds are not seeing the blades at all. While vultures did not occur on our site many raptors that forage for terrestrial prey exhibit similarly wide blind areas due to skull morphology (Potier et al. 2018) and were common on site. For such species (eagles, harriers, kestrels, Elanus kites) none were killed by the blades once patterned, despite falling victim before patterning. The approximate 88% reduction in fatalities for this group as a whole at Hopefield suggests that skull morphology has little influence on susceptibility to blade strikes, but it cannot be ruled out for vultures until similar experiments occur in areas within their range.

A second hypothesis has arisen from the finding that many raptors have very poor achromatic contrast abilities up to ten-fold lower than human vision (Potier et al 2018, Blary et al 2024). This may adequately explain a raptor’s inability to detect white turbine blades against a pale background (even when flying directly at a set of blades: Simmons and Martins 2024) and explain why increasing the contrast in the lab (McIsaac 2001) or in the field (May et al. 2020, this study) makes a high contrast blade more conspicuous to raptors. The patterning itself may have the added advantage of rendering the blade intrinsically more conspicuous (i.e. along the blade itself) and against a bright background.

We inadvertently tested the use of an alternative colour (Signal Red) at Hopefield due to constraints imposed by the South African Civil Aviation Authority on the use of black paint. This is both a useful result (for other areas where heating of black blades may compromise blade integrity or not permitted e.g. Santander et al. 2026) and interesting for exploring what cues raptors may be using (colour or contrast).

We must ask why red patterns worked well in this experiment when traditionally, avian research indicates that the retina responds best to high contrast combinations especially at low light levels (Potier et al. 2018, Martin and Banks 2023, Blary et al 2024). That is, vision experts promote the use of striped black and white turbine blades as it provides the highest achromatic contrast (4% and 85% reflectance) and better than Signal Red and Pure White (with 8% and 85% reflectance). There are two probable answers: birds respond better to patterns than solid colours in the lab (McIsaac 2001, Blary et al. 2026 Klein Heerenbrink et al 2025, Hancock et al. 2026) but red may have an added advantage of acting as an aposematic warning colour. Recent lab experiments with red alone, black alone and red/orange and black stripes together on turbine blades (Hancock et al. 2026) indicated that a European passerine responded behaviourally more emphatically to the aposematic colours than red or black alone. Red and orange are colours have evolved in a vast array of aposematically coloured species (e.g. butterflies, bees, beetles, birds, amphibians, reptiles and mammals) and are easily remembered and elicit an aversion even in naïve birds (Halpin et al. 2020, Hancock et al. 2026). Therefore, we posit that red patterns used in this experiment may be more effective than black patterns because it introduces a behavioural deterrent, over and above it simply being more visible to birds. Further experiments in the lab and field are required to tease apart the merits of behavioural aversion vs retinal responses in the optimal design and colour to reduce avian impacts.

Despite the small samples in both countries (four experimental turbines each in Norway and South Africa) both studies found a positive effect on a range of different species and one particularly vulnerable group - the raptors (Thaxter et al. 2017, Watson et al. 2018, Ralston-Paton 2025). At Smøla, follow-up research after the main experimental period (Stokke et al. 2024) revealed a 100% reduction in eagle fatalities at the black blade turbines despite no reduction in the average number of eagles killed elsewhere in the facility. In Hopefield, with 14 species of raptor present, and at least six species breeding in or near the facility (one eagle, one harrier, one kite, one buzzard and two falcon species) 10 species have been killed at the turbines. Raptors accounted for 33% of all fatalities prior to patterning and 21% after patterning. Despite raptors continuing to be killed at Hopefield’s control turbines (8 deaths) only one raptor (a *Milvus* kite) was killed at the experimental turbines in 32 months, a decrease of ~86% over the seven killed prior to treatment. The most vulnerable of those to population instability, the Endangered Black Harriers (Cervantes et al. 2022), suffered no further deaths at the patterned blades. Thus, both studies show that increasing blade contrast is effective for one of the most impacted groups - the raptors.

Two of the concerns that this study does not directly address are (i) whether the patterning would be effective in low light conditions – because the achromatic contrast provided by red and white (in this study reflectance 8% vs 85%) is likely to be lower than black and white (Martin and Banks 2023) and (ii) its effectiveness in reducing collisions for the most frequent red data victims at South African wind farms such as Verreaux’s Eagles *Aquila verreauxii* and Cape Vulture *Gyps coprotheres* (Ralston-Paton 2025).

The poor light conditions were partially tested when it emerged that Hopefield experiences 110 fog days per year (Olivier 2004), implying that blade patterning is effective at least in foggy conditions. However other reduced visibility conditions and effectiveness at dawn, dusk and at night remain unexplored. In addition, we must await field tests from other wind farms with vultures to complement the lab studies suggesting patterns similar to those here are effective for Gyps vultures (Blary et al. 2026). Similarly, despite the presence of four eagle species around Hopefield the Verreaux’s Eagle, South Africa’s most impacted eagle (Ralston-Paton 2025), was not present and blade patterning trials within their range would prove insightful.

We must also acknowledge that choosing high fatality turbines, rather than selecting turbines randomly for treatment may skew our results. In a practical sense this may be good conservation practice, but the highest fatality turbines may result in the highest reductions because they experienced higher fatality rates. We justified this in the knowledge that fatalities are rare events (REWI 2025, Ralston-Paton 2025) and achieving significant results within the financial constraint imposed by carcass searching year after year sets an upper limit on the time available for such experiments.

We encourage future trials to pattern all three blades as used in most lab experiments to date (McIsaac 2001, Blary et al 2026, Klein Heerenbrink et al. 2025, Hancock et al. 2026), and not one blade, as promoted by Hodos (2003). Motion smear seems unlikely to be behind the high fatality rate attributed to wind turbines because modern turbines do not spin fast enough to produce such an effect (H. McIsaac, C. Blary, M. Klein Heerenbrink pers comm.).

### Blade patterning relative to other mitigation measures

The magnitude of the decline in numbers of fatalities at the patterned turbines at Hopefield ranks it among the most successful operational-phase mitigations thus far tested in the field. According to independent reviews (McClure et al. 2021, Garcia-Rosa and Tande 2023) of different forms of passive and active mitigation (i.e. automated shut down on demand, feathering of blades, auditory deterrents, blade painting and habitat management), the latter, tilling of soil near turbines to reduce prey populations for raptors (Pescardo et al. 2019), achieved the highest success in reducing raptor deaths (86%) before and after implementation. This was followed by automated shut down for two eagle species (82% success: McClure et al. 2021). The research also identified drawbacks to some of the mitigations including habitat loss, expense and unnecessary shutdowns (false positives) for the technological mitigations reviewed. The beauty of the passive red stripes or black blade approach is that once the mitigation is in place, there are no post-application costs, and for wind farms implementing blade patterning at source, there are no extra costs associated with painting blades in situ, in windy conditions. The blades can be ordered pre-patterned from the factory as necessary.

## Conclusions and recommendations

We conclude that the patterning of turbine blades at the species-rich Umoya Wind farm in South Africa with two broad signal red stripes significantly reduced avian fatalities by a median 83% at the treated turbines, supporting the results of the first study in Norway. That Signal Red paint (with reflectance of 8%) showed comparable results to the black-painted blades in Norway (with ~4% reflectance) suggests that as long as high achromatic contrast paints are used on white blades, birds are more likely to see and avoid operational turbines with this mitigation. We additionally suggest that a patterned design may be more visible to birds than one solid design given the magnitude of the effect of blade patterning at the Umoya wind farm, the lack of a positive result at the Dutch black-blade experiment (Klop et al. 2025), and the increased detection of striped patterns for four species in laboratory experiments (McIsaac 2001, Blary et al. 2026). That red as an aposematic signal may add a behavioural deterrent for birds (Hancock et al. 2026) gives a further dimension to red-blade patterning as a mitigation.

We call for other wind farms in other high biodiversity regions with different suites of collision-prone species to trial suitably patterned blades to reduce fatalities. This will determine its efficacy across multiple species and environments, nocturnally and diurnally.

## Acknowledgements

Umoya Energy (S. Abrahams, R. Hammond and N. Luttig) were instrumental in prising open the bureaucratic doors permitting the patterned blades to be trialled, and L. Strohl, N.Mojela and N. Nkabiti of SACAA enabled CAA approval. We are grateful to all Umoya carcass searchers for their long hours and commitment over the years. S. Ralston-Paton (Birdlife South Africa), S. Taylor (South African Wind Energy Association), JLB Simmons and J. Suri (University of Cape Town), A. Pearson (Arcus) all assisted in the completion of this project. We thank them and Drs R. May, B. Iuell, R. Diehl, S. Felton, S. Childs, T. Katzner, M. Klein Heerenbrink, E Klop, J. Kleyheeg Hartman, E. Sorondo, Professors G. Martin, G. Taylor and especially C. Blary and S. Potier for many fruitful discussions and reviews.

---

**Supplementary Table 1.**
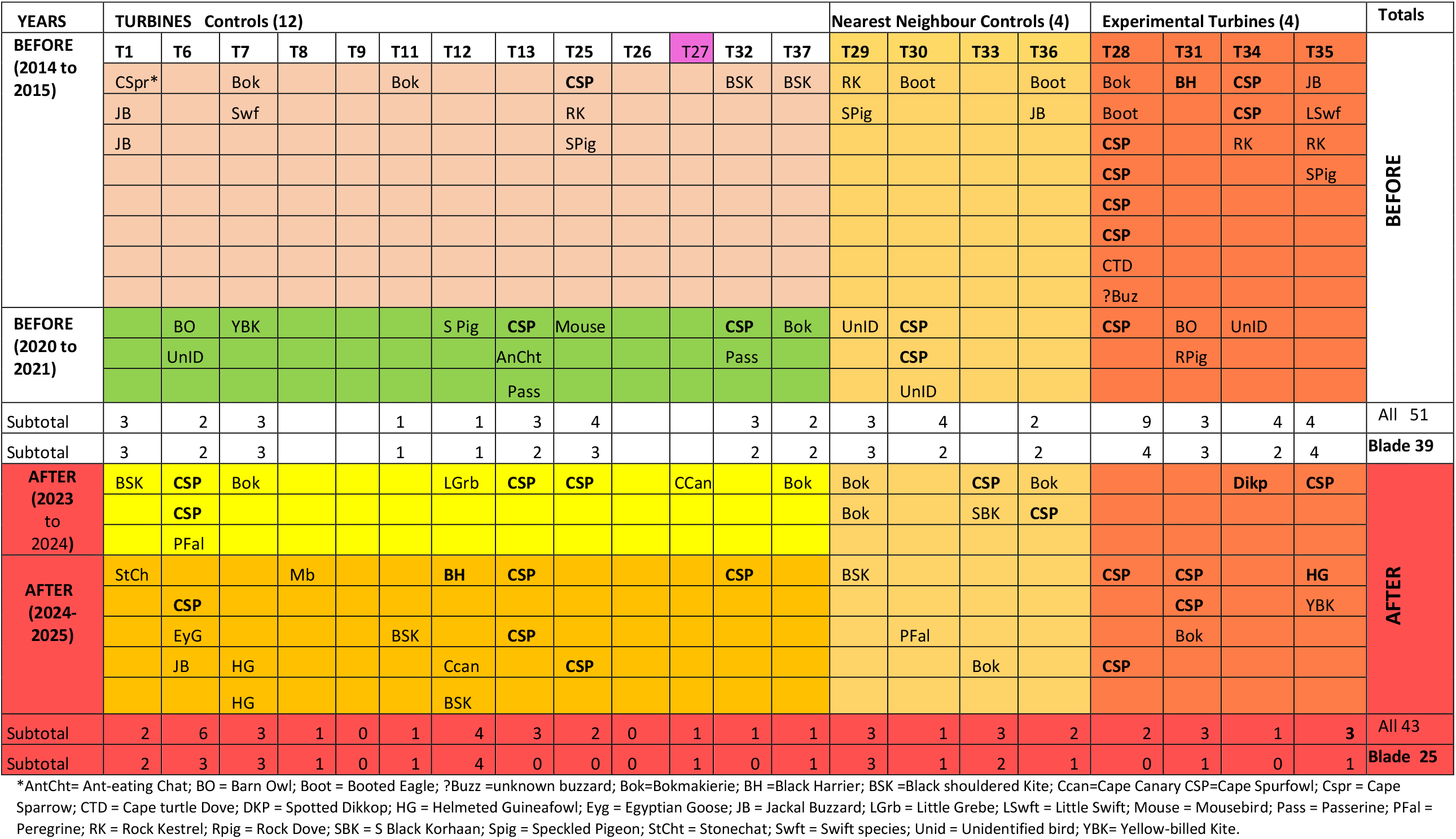
All fatalities across all (20) turbines and all study years Before (2014/15, 2020/21) and After (2023/24/25) treatment. Nearest neighbour controls are distinguished from other controls as they are used separately to test for an effect of blade patterning at the Experimental turbines. Two subtotal are given for both periods (before and after): the first includes tower victims (like Cape Spurfowl [CSP]) and the second omits them to give just the blade strike victims. Turbine T27 was substituted for T26 while T26 was offline.

## Notes

### Competing Interest Statement

The authors have declared no competing interest.

